# The UCSC Repeat Browser allows discovery and visualization of evolutionary conflict across repeat families

**DOI:** 10.1101/429613

**Authors:** Jason D. Fernandes, Armando Zamudio-Hurtado, W. James Kent, David Haussler, Sofie R. Salama, Maximilian Haeussler

## Abstract

**Background:** Nearly half the human genome consists of repeat elements, most of which are retrotransposons, and many of these sequences play important biological roles. However repeat elements pose several unique challenges to current bioinformatic analyses and visualization tools, as short repeat sequences can map to multiple genomic loci resulting in their misclassification and misinterpretation. In fact, sequence data mapping to repeat elements are often discarded from analysis pipelines. Therefore, there is a continued need for standardized tools and techniques to interpret genomic data of repeats.

**Results:** We present the UCSC Repeat Browser, which consists of a complete set of human repeat reference sequences derived from the gold standard repeat database RepeatMasker. The UCSC Repeat Browser contains mapped annotations from the human genome to these references, and presents all of them as a comprehensive interface to facilitate work with repetitive elements. Furthermore, it provides processed tracks of multiple publicly available datasets of biological interest to the repeat community, including ChIP-SEQ datasets for KRAB Zinc Finger Proteins (KZNFs) – a family of proteins known to bind and repress certain classes of repeats. Here we show how the UCSC Repeat Browser in combination with these datasets, as well as RepeatMasker annotations in several non-human primates, can be used to trace the independent trajectories of species-specific evolutionary conflicts.

**Conclusions:** The UCSC Repeat Browser allows easy and intuitive visualization of genomic data on consensus repeat elements, circumventing the problem of multi-mapping, in which sequencing reads of repeat elements map to multiple locations on the human genome. By developing a reference consensus, multiple datasets and annotation tracks can easily be overlaid to reveal complex evolutionary histories of repeats in a single interactive window. Specifically, we use this approach to retrace the history of several primate specific LINE-1 families across apes, and discover several species-specific routes of evolution that correlate with the emergence and binding of KZNFs.

## INTRODUCTION

Transposable elements are significant drivers of eukaryotic genome evolution. In humans and other primates, transposons constitute nearly half the genome; the majority of these repeat elements are retrotransposons, although some DNA transposons are also present. Despite the high repeat content of the human genome, many genomic analyses struggle to deal with these regions as sequencing reads can often be assigned nearly equally well to multiple regions in the genome. Masking or filtering these reads is often considered a “conservative” approach in that it avoids mis-assigning the genomic location of a read, but it prevents the discovery of important biology occurring at repeat elements^1^. Indeed, many repeats already have established roles in important biological processes, complex behavioral phenotypes, and disease^2–5^.

One of the major challenges in proper repeat-analysis is establishing a set of standardized sequences, nomenclature and annotation sets that can be universally understood by the scientific community. The most commonly used databases and tools to study repeats are Repbase^6^ and RepeatMasker^7^. Repbase began as a hand-curated list in 1992 of 53 prototypic repeat sequences identified in the human genome^8^. By 2015, it contained more than 38,000 sequences in 134 species^6^, making curation and comprehension of each repeat family a daunting challenge. RepeatMasker is a program that screens DNA (e.g. a newly sequenced genome) for repeat elements. RepeatMasker utilizes a specialized version of RepBase (RepBase RepeatMasker Edition) as input to identify repeats within a genome. RepeatMasker’s final output also represents additional optimizations (e.g. building full length repeat elements from smaller subparts, generalization (grouping together) of similar elements, and specialization (using information about repeat structure)) designed to improve the speed and quality of repeat detection (Figure 1A).

**FIGURE 1:**
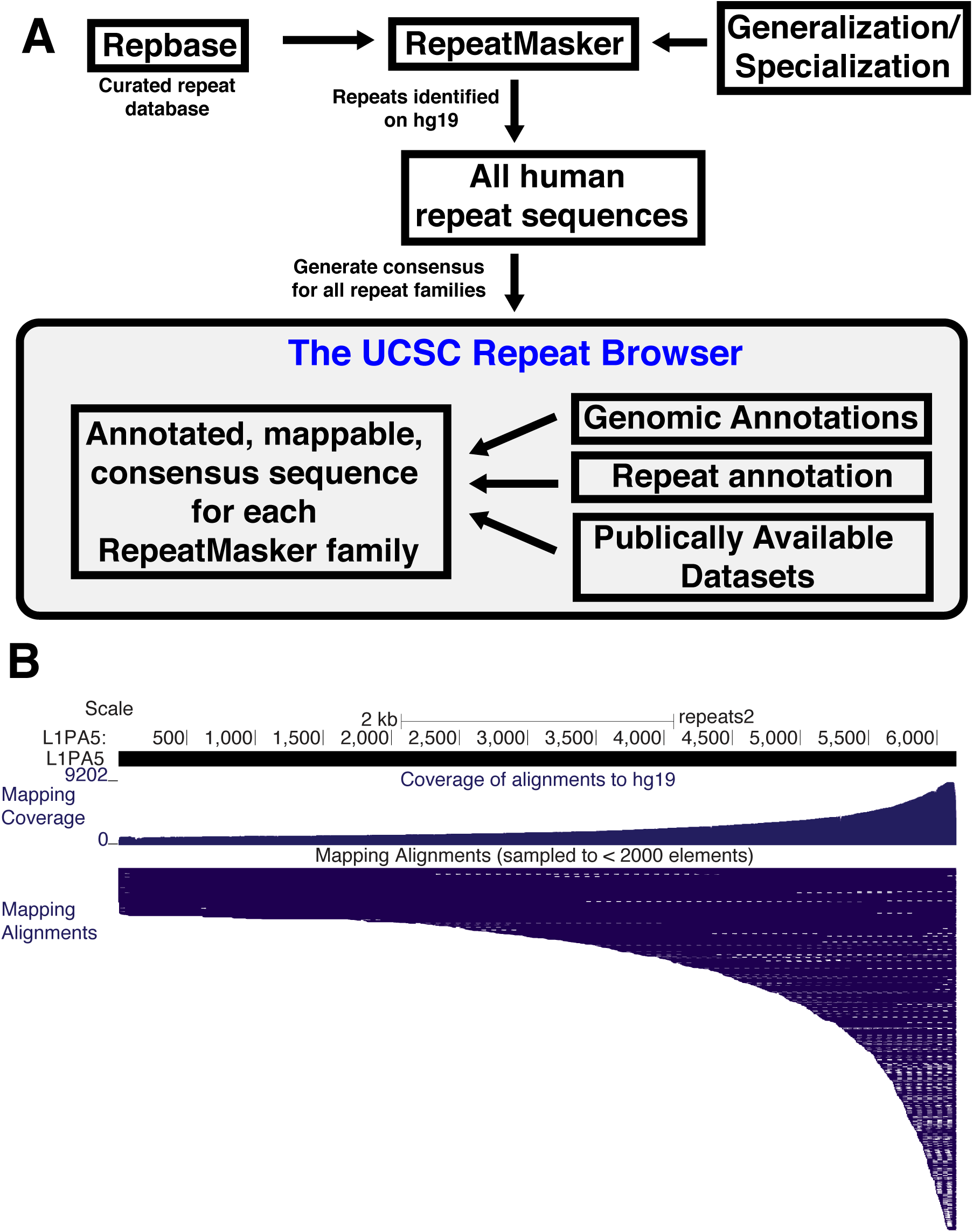
Building the UCSC Repeat Browser. A) Workflow for building the UCSC Repeat Browser. Repeat annotations and sequences are taken directly from RepeatMasker tracks across the human genome and used to build reference consensus sequences for every repeat family. Existing genomic annotations are then mapped to these consensuses. B) Mapping of all individual L1PA5 instances to the consensus. A majority of L1PA5 sequences in the human genome only contain the 3’ end as evidenced by the coverage per base (mapping coverage) and alignments of individual instances (mapping alignments).

Although a variety of tools and methods already exist to study repeats^9^, tools to dynamically visualize genomic data and interact with existing annotation sets on repeats (e.g. protein coding regions, conservation with other sequences and the list of matches in the genome) are currently underdeveloped. Generating and mapping to a consensus version of individual repeats has proven successful in illustrating novel biological features of transposon insertions, but has largely been limited to static visualizations on targeted elements of interest and specific families of these repeats^10, 11^.

Here we present the UCSC Repeat Browser, which simplifies analysis of genomic data on repeats by providing automatically generated consensus sequences for all human repeat element classifications within RepeatMasker. The Repeat Browser overlays a precomputed set of comprehensive annotations in an interactive genomic browser environment (Figure 1). Further, we demonstrate the utility of the Repeat Browser in uncovering and illustrating evolutionary conflict between a primate specific class of retrotransposons and their repressors.

## IMPLEMENTATION

### Generating Reference Sequences for Human Repeats

We first generated consensus reference sequences for each repeat family listed in the RepeatMasker annotation of the human genome (hg19). To do so, we downloaded all nucleotide sequences and their annotations in the RepeatMasker annotation track on the UCSC Human Genome Browser (hg19). We observed that extremely long repeats tended to represent recombination or misannotation events and therefore removed the longest 2% of sequences in all classes. We then aligned the 50 longest remaining sequences of each class, as this produced a tractable number of sequences that allowed manual inspection of each alignment, and because insertions relative to the consensus are otherwise invisible when plotted on shorter sequences. For each repeat family, these fifty sequences were realigned with MUSCLE^12^ to create a consensus sequence. Each of these consensus sequences was then stored as a “reference” in the Repeat Browser in a manner analogous to a single chromosome on the UCSC Human Genome Browser^13, 14^. Each alignment is provided as a link in a “consensus alignment” track for additional visual inspection by the user.

### Annotation of each repeat class

For each repeat family, the consensus was mapped back to all of its repeats with BLAT^15^. From this process, we generated a coverage plot illustrating the relative representation of the consensus from each genomic instance (Figure 1B). For example, the primate-specific LINE-1 sub-family, L1PA5, shows the expected distribution: most of the individual L1PA5 instances, are short 3’ truncations, meaning that most genomic loci annotated as L1PA5 do not contain the 5’ portion. Therefore the 3’ end of the consensus is found relatively more often across the human genome (Figure 1B). We also ran Tandem Repeats Finder^16^ and the EMBOSS ORF finder^17^ on these consensus sequences in order to automatically annotate each consensus. We similarly aligned the RepeatMasker Peptide Library^18^ with BLAST^19^ and each of the original genomic sequences with BLAT^15^ to each consensus.

Our alignment of individual repeat elements in the genome to their respective consensus sequence allows us to map any genome annotation to the genome consensus sequence, a process more generally known as “lifting”. In this way, human genes that contain repeat sequence (as annotated by GENCODE^20^ and UCSC genes^21^) were “lifted” to each consensus sequences (Table 1). Figure 2 shows the results for L1PA5 elements. As expected, L1PA5 sequences that have been incorporated into protein coding genes tend to derive from the untranslated regions (UTRs) of the repeats and have incorporated into the UTRs of the protein coding genes. Finally, although the Repeat Browser consensus sequences are built from hg19 RepeatMasker annotations, we also generated mappings of each consensus to each corresponding repeat instance in hg38. The result of these procedures produces a fully annotated and interactive consensus sequence that requires minimal prior knowledge of the genomic organization of the repeat being analyzed and at the same time allows lifting of any genome annotation from either hg19 or hg38.

**FIGURE 2:**
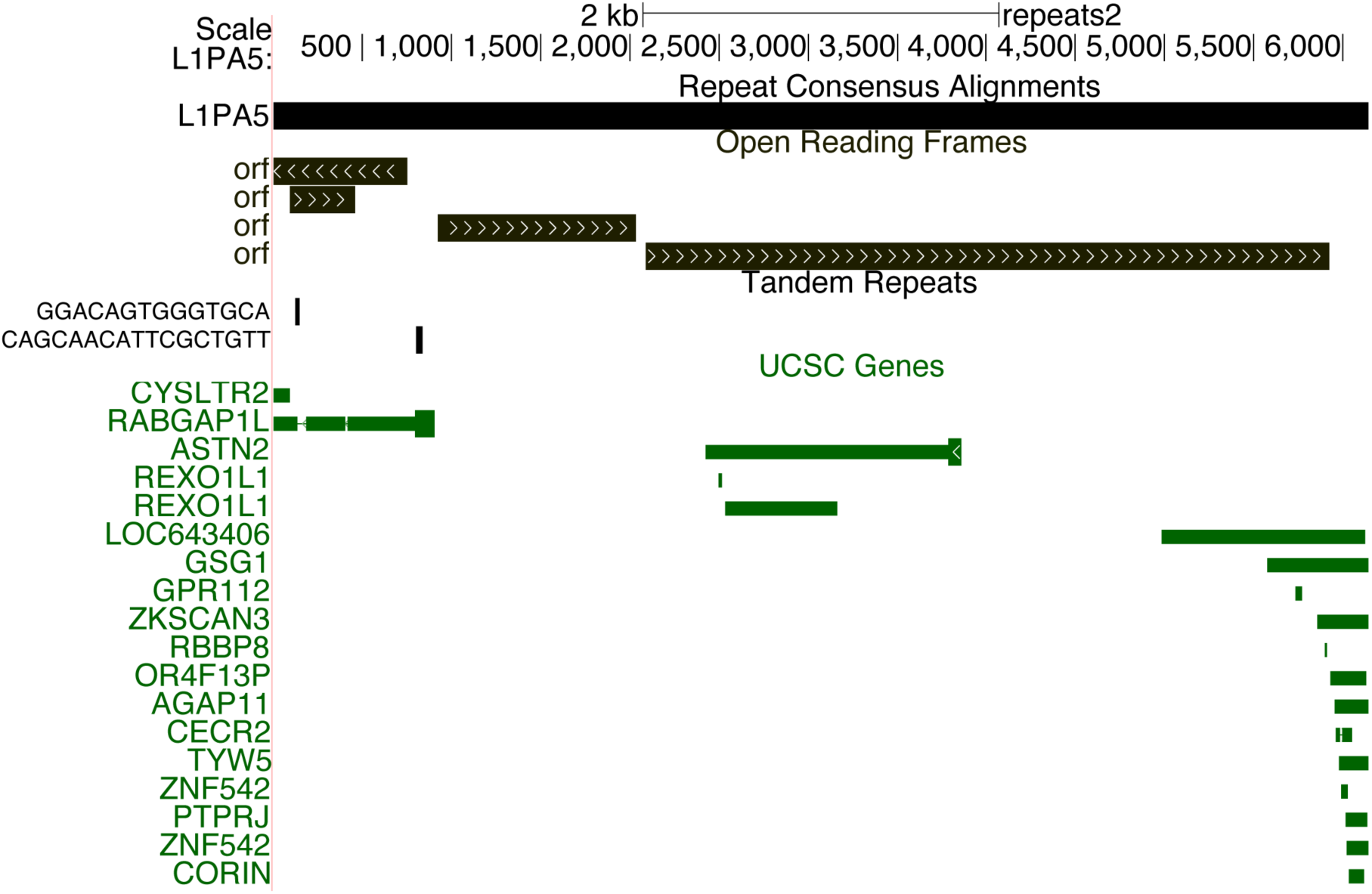
Mapping of existing annotations and detection of repeat features. Annotation sets (e.g. UCSC Genes) that intersect RepeatMasker annotations were lifted from hg19 to the Repeat Browser consensuses. Shown here are all genes that contain L1PA5 sequence as well as ORFs (detected by EMBOSS getorf) and tandem sequence repeats detected within the L1PA5 consensus detected by Tandem Repeat Finder.

**Table 1.** List of Tracks available on the Repeat Browser.

| Track | Description |
| --- | --- |
| Mapping Alignments | Alignments of each individual repeat instance in hg19 back to the Repeat Browser consensus. |
| Mapping Coverage | A coverage plot for the mapping alignments. |
| Conserved Elements | Highly conserved genomic sequences in vertebrates, placental mammals and primates lifted to the Repeat Browser. |
| RepeatMasker Proteins | Protein products of the repeat element as annotated in RepeatMasker records. |
| ORFs | Predicted ORFs |
| Other Cons Aln | Alignment of all other Repeat Browser Consensuses against the currently viewed consensus. |
| Repeat Consensus Alignments | Alignment of all repeats from the RepBase RepeatMasker Libraries |
| Tandem Repeats | Detected tandem sequence repeats within the consensus full-length repeat elements. |
| ENCODE Tracks | DNase mapping, histone marks and TF ChIP-SEQ from ENCODE lifted to the Repeat Browser. |
| KZNF Tracks (Imbeault/Trono 2017 & Schmittges/Hughes 2016) | Lifting of reprocessed data from large KZNF ChIP-SEQ screens. |
| TF Differentiation Data (Tsankov 2014) | Lifting of large scale ChIP-SEQ dataset from differentiation time course across multiple cell types. |
| Stem Cell State Data (Theunissen 2016) | Lifting of reprocessed data from primed and naïve human pluripotent stem cells. |

### Mapping of Existing Genomic Datasets

We also mapped genomic loci bound by histone-modifying enzymes from ENCODE datasets^22^ as well as large-scale ChIP-SEQ collections KRAB Zinc Finger Proteins (KZNFs) ^23, 24^ to the Repeat Browser. KZNFs are particularly compelling factors as they engage in evolutionary “arms races” in which KZNFs evolve unique DNA binding properties to bind and repress retrotransposons^10, 25^. These retrotransposons then accumulate mutations that allow evasion of KZNF-mediated repression^10^. In order to map this ChIP-SEQ data to the Repeat Browser, we first downloaded raw ChIP-SEQ reads from the Sequence Read Archive (SRA)^26^, mapped them to the reference genome (hg19) using bowtie2^27^ and called peaks using macs2^28^ (Figure 3A). After this standard genomic mapping and peak calling, we then took the peaks of these these DNA-binding summits that overlapped a repeat element as annotated in the RepeatMasker track, extended them by 5 nt in both directions, and used BLAT to map them to the appropriate (as determined by RepeatMasker annotation) Repeat Browser consensus sequence. In essence, this approach leverages each repeat instance as a technical replicate, with the mapping to the consensus representing a combination of many genomic “replicates’ (Figure 3A) of DNA binding summits called on individual instances of a repeat family that individually produce a noisy set of mappings; however hundreds of them combined yield a clear overall signal, better identifying the actual binding site. We call this “summit of summits” (obtained by combining the summits on individual transposon instances into a single summit on the Repeat Browser consensus) the “meta-summit”. In order to determine these “meta-summits”, we ran our peak caller (macs2) on the repeat consensus to generate a list of “meta-summits” which represent the most likely location of the DNA binding site for a specific DNA-binding factor. We then generated a track which summarizes these meta-peaks for each consensus sequence allowing easy and quick determination of factors with correlated binding patterns (Figure 3B; visualized on Human Endogenous Retrovirus H (HERV-H)).

**FIGURE 3:**
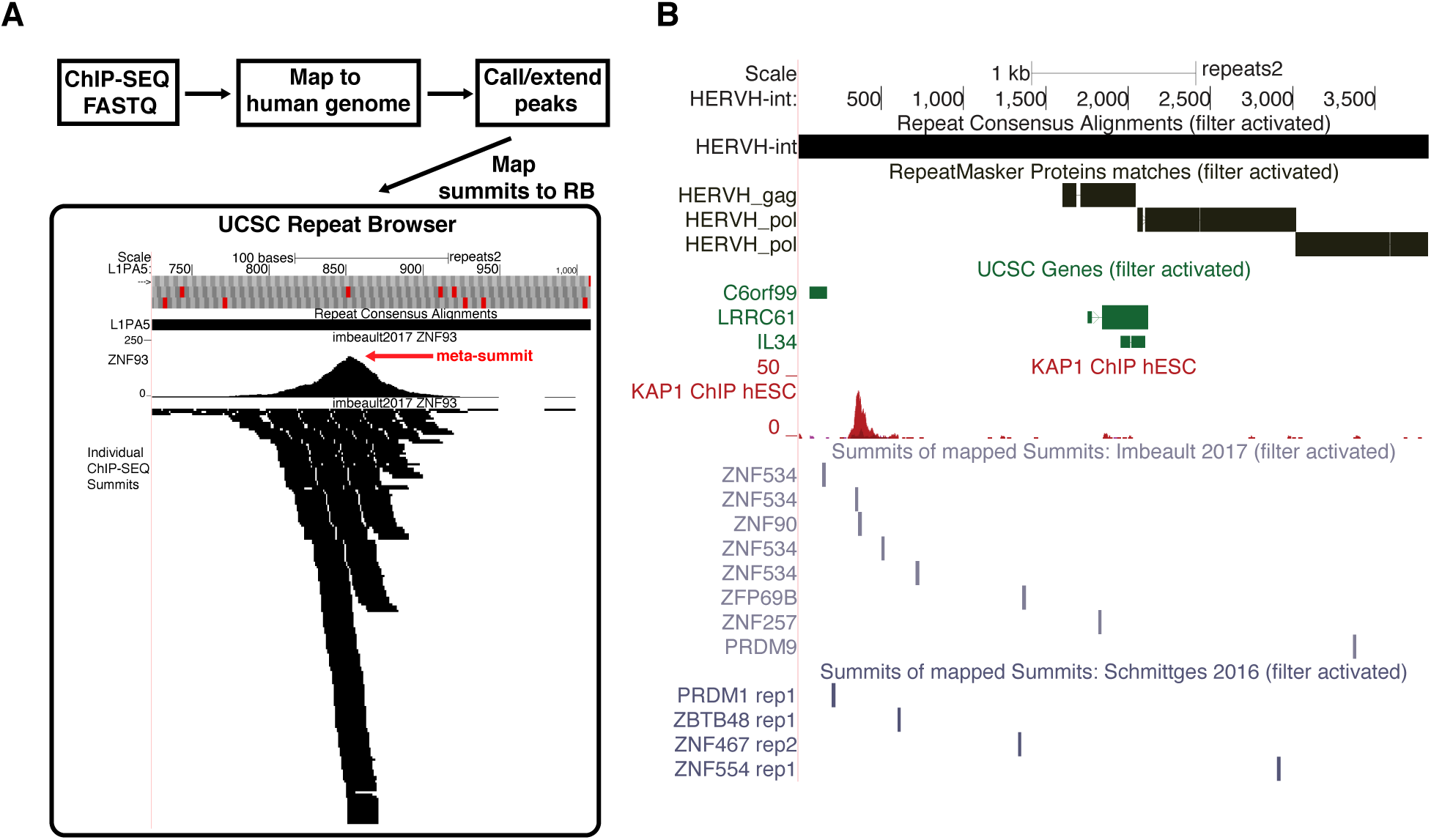
Mapping of KZNF ChIP-SEQ data to the UCSC Repeat Browser. A) Workflow for analyzing KZNF ChIP-SEQ. Data from existing collections was downloaded from SRA, analyzed via standard ChIP-SEQ workflows and the resulting summits mapped back to the RB for analysis. Mapping of individual summits produces a “meta-summit” (red arrow) that can be used for downstream analysis and which is stored separately in another annotation track. B) Example of a repeat family, HERVH-int (a primate endogenous retrovirus) with lifted annotations and datasets. Shown are tracks of annotated ORFs, gene overlaps, Kap1 ChIP-SEQ coverage and KZNF meta-summits.

## RESULTS

### Comparative Analysis of L1PA elements

In order to demonstrate the power of the UCSC Repeat Browser, we studied the evolution of recent L1PA families. The L1PA lineage is a group of LINE-1 retrotransposon families specific to primates. These elements are fully autonomous, and encode proteins (ORF1 and ORF2) responsible for reverse transcription and re-integration of the retrotransposon. L1PA families evolve in bursts; higher numbers (e.g. L1PA17) indicate ancient evolutionary origins, as evidenced by shared copies across species (Fig 4A). Lower numbers indicate more recent activity and are derived from the older, higher number families (note L1PA1 is also known as L1HS, human-specific)^29^. Although this nomenclature generally corresponds to speciation events on the phylogenetic tree of the hosts of L1PA retrotransposons, many families had overlapping periods of activity meaning that the correspondence is not exact (e.g. it is possible that a few L1PA3 instances are present in gibbon, despite their major burst of activity on the human lineage occurring after the human-gibbon divergence)^30^.

**FIGURE 4:**
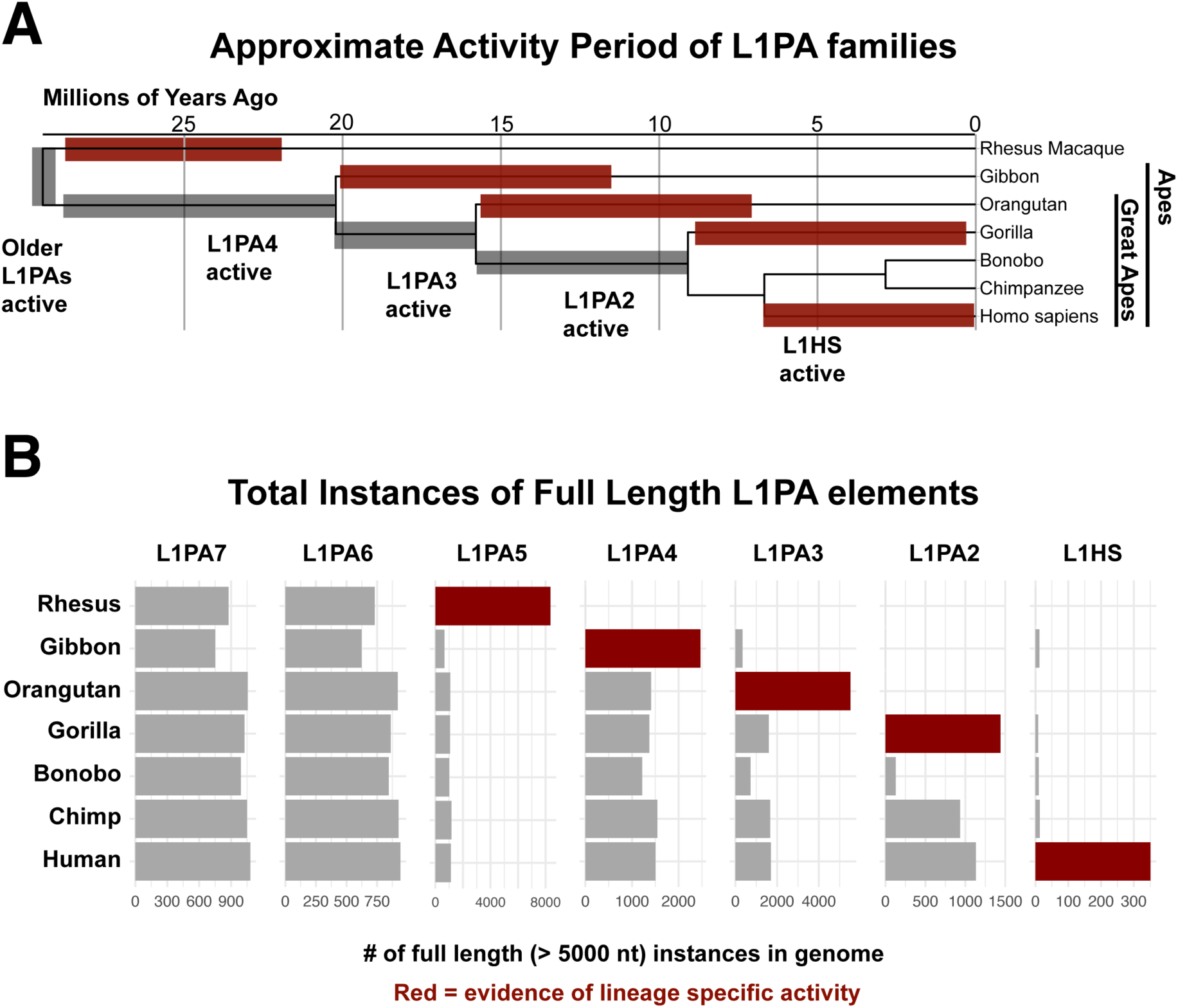
Comparative analysis of L1PA elements. A) Phylogeny and nomenclature of L1PA elements. Older elements have higher numbers and families can expand in a manner that will be conserved between species (grey) or lineage-specific (red). B) Counts of full length L1PA instances extracted from UCSC Repeat Masker tracks. Note for Rhesus (rheMac10), L1PA5 counts represent a sum of rhesus-specific elements (labeled as L1PA5 in RepBase, L1_RS* by RepeatMasker). Families in red expand greatly compared to families in grey, providing evidence of lineage-specific expansion.

### Comparison of Primate Repeat Elements Reveals a Large Number of Gibbon Specific L1PA4 Elements

In order to trace the evolution of L1PAs in different species, we downloaded the complete sequences for every L1PA7 and younger L1PA family, as annotated in their UCSC Genome Browser RepeatMasker tracks, in rhesus macaque (rheMac10), gibbon (nomLeu3), orangutan (ponAbe3), chimp (panTro6), gorilla (gorGor5), bonobo (panPan2) and human (hg38). We further restricted our analysis to only full-length elements by filtering out elements less than 5000 nucleotides in length.

As expected, the number of elements in older families were largely similar amongst all species that shared a common ancestor when the retrotransposon was active: for instance, L1PA7, active prior to the emergence of the last common ancestor of all primates in this study, was found at a relatively constant amount in all genomes (Figure 4B). On the other hand, human specific elements were found only (barring a few likely mis-annotations) in the human genome. Curiously, in certain species (gibbon, orangutan and gorilla) instances of retrotransposon families that were active near their divergence from human, were present in much greater copy number (Figure 4B). Specifically, the number of L1PA4 elements was greater in gibbon then all other apes, while a similar pattern was seen for L1PA3 and orangutan, and L1PA2 and gorilla. These results are consistent with these primates having lineage specific expansion of these elements in a manner distinct from humans. Notably, bonobos had a markedly lower number of L1PA2 elements which may indicate stronger repression of these elements by a species-specific factor; however, the bonobo assembly was one of the older, short-read primate assemblies used in this study, and therefore the lack of L1PA2 elements may simply reflect greater difficulty in resolving these regions in the genome assembly. Note also that the UCSC track for rheMac10 contains no annotated instances of L1PA5, but this simply reflects the fact that RepeatMasker taxonomy splits the L1PA5 family into L1_RS families that are rhesus-specific compared to the other primates in this study^31^.

### All apes display evidence of ZNF93 evasion in the 5’UTR of L1PA

In order to examine the selection pressures that might explain species-specific expansion and restriction of L1PA elements, we combined our primate L1PA analysis with the ChIP-SEQ data of KRAB Zinc Finger Proteins (KZNFs) on the Repeat Browser^23, 32^. KZNFs rapidly evolve in order to directly target retrotransposons and initiate transcriptional silencing of these elements. We previously demonstrated that a 129bp deletion occurred and fixed in the L1PA3 subfamily (and subsequent lineages of L1PA) in order to evade repression mediated by ZNF93. In order to discover additional cases where a retrotransposon may have deleted a portion of itself to escape KZNF-mediated repression, we analyzed L1 sequences with the following characteristics: 1) deletion events proximal to KZNF binding sites, and 2) increasing number of retrotransposon instances with that deletion (demonstrating increased retrotransposon activity). Comparisons of these events across primate species, provides evidence for unique, species-specific mechanisms of escape.

In order to look for these signatures of L1PA families escaping repression, we used BLAT to align each individual full-length (>5000 nt) primate L1PA of the same class instance to the human Repeat Browser consensus from the primate genomes under study. We then generated coverage tracks of these full-length elements mapped to the human consensus for each species and each L1PA family. The ZNF93-mediated deletion is clearly visible as evidenced by a massive drop in coverage in the 129-bp region in human L1PA3 instances (Figure 5A). This same drop in coverage is found in all great apes (orangutan, gorilla, bonobo, chimp, and human) confirming that this event occurred in a common ancestor. Notably a small number (∼300) of L1PA3 elements were identified in gibbon; however these elements display a different drop in coverage (20 bp long) near the ZNF93 binding site, The majority of these gibbon “L1PA3” instances do not lift to the human genome (or lift to older L1PA5 and L1PA4 elements) suggesting they are mis-annotations or gibbon-specific L1PA expansions. Therefore, we examined gibbon L1PA4 elements on the Repeat Browser and found that the small 20 bp deletion - at the base of the ZNF93 peak – first occurred in Gibbon L1PA4 elements (Figure 5B), after the human-gibbon divergence (since humans and other great apes do not have this deletion), and likely gave rise to gibbon-specific L1PAs. Elements with this 20-bp deletion were likely able to evade ZNF93, and may also hold a selective advantage over more drastic 129 bp L1PA3 deletions. Indeed, elegant work from the Moran lab has recently shown that the 129bp deletion in human L1PA3 elements alters L1PA splicing in a manner that can generate defective spliced integrated retrotransposed elements (SpIREs)^33^: the smaller deletion found in gibbons may avoid generating these intermediates. Additionally, gibbon L1PA4 elements also experience a smaller coverage drop (typically near the ZNF765 binding site (Figure 5B). Coverage drops in this area are found predominantly in L1PA4 instances with the ZNF93 binding site already deleted, indicating that this deletion (and the presumed escape from ZNF765 control) occurred after escape from ZNF93 control (Figure 5C).

**FIGURE 5:**
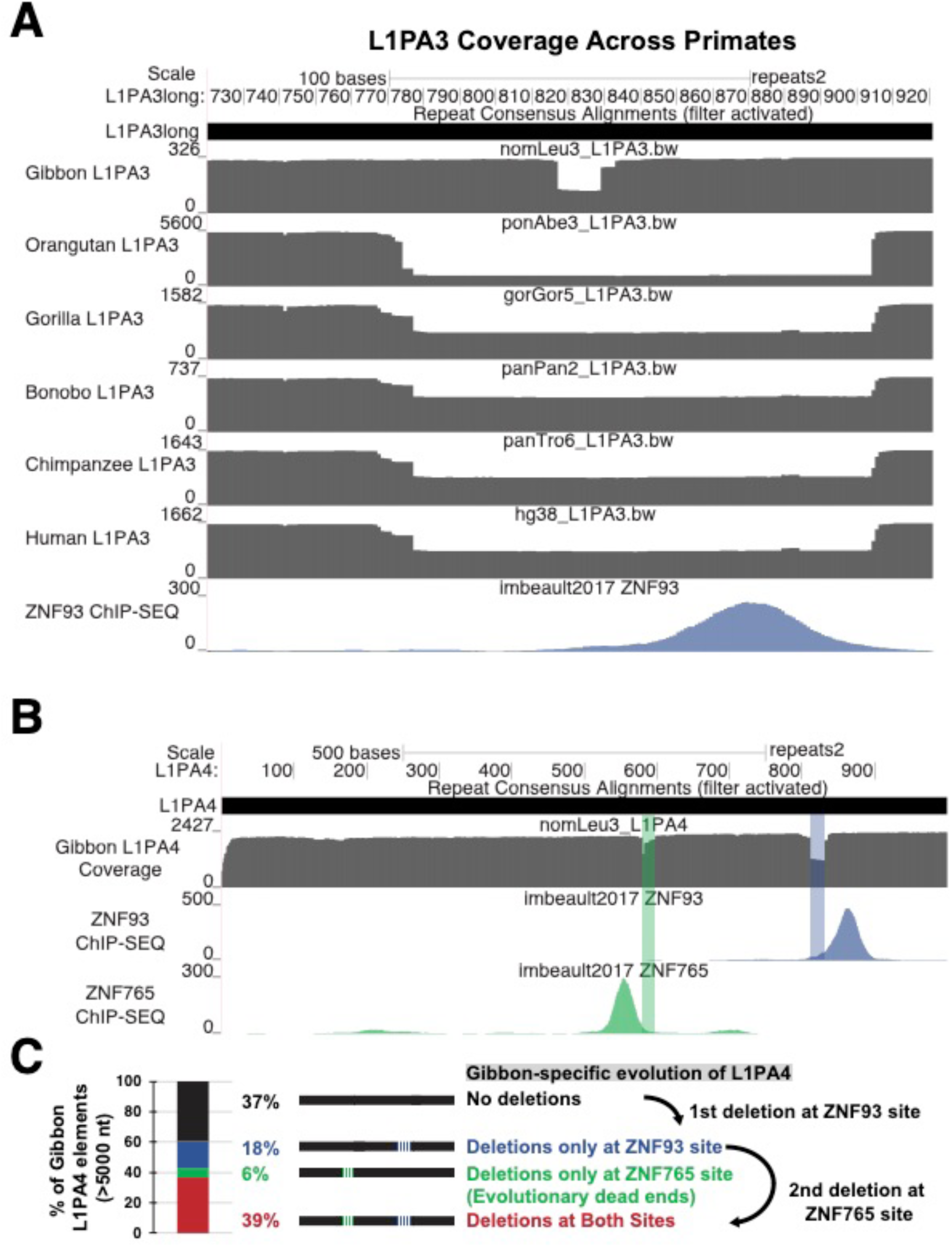
Comparative analysis of L1PA3 & 4 elements in apes and great apes. A) Coverage tracks for all full length ape L1PA3 elements mapped to the human consensus. Gibbons have few L1PA3 elements that are likely misannotated L1PA4 elements and a unique deletion in the ZNF93 binding (blue) region. All great apes (all shown except gibbon) exhibit a shared deletion, evidenced by a coverage drop over 129 bp. B) Coverage map of gibbon L1PA4 elements demonstrates a different path of ZNF93 evasion (20 bp deletion) as well as a second region spanning 22 bp near the major ZNF765 binding site (green). (Below) Analysis of mutational patterns in gibbon demonstrates that the 20 bp ZNF93-associated deletion likely occurred first in gibbon L1PA4 as most L1PA4s with ZNF765-associated deletions also contain a ZNF93-associated deletion.

### Novel Orangutan-Specific Deletions are Visible on the UCSC Repeat Browser

L1PA3 elements display an increased copy number in the orangutan genome, suggesting that these elements also had a lineage specific expansion, driven by escape from KZNFs or other restriction factors. Aligning of orangutan L1PA3 elements on the Repeat Browser L1PA3 consensus displayed a clear 11 bp deletion ∼230 bp into the 5’ UTR that is not present in human, chimp or bonobo elements (Fig 6A). However, analysis of existing KZNF ChIP-SEQ data, shows no specific factor that clearly correlates with this deletion. We may simply lack ChIP-SEQ data for the appropriate factor (including the possibility that the KZNF driving these changes evolved specifically within the orangutan lineage) to explain the evolutionary pattern seen in these orangutan-specific elements; alternatively, this mutation might alter some other aspect of L1PA fitness (e.g. splicing). Regardless, L1PA3 elements with this deletion were highly successful in spreading throughout the orangutan genome. Furthermore, L1PA3 instances with deletions in this region also harbor the 129 bp ZNF93 deletion, suggesting that this 11 bp deletion occurred after orangutan L1PA3 elements escaped ZNF93 control (Fig 6B).

**FIGURE 6:**
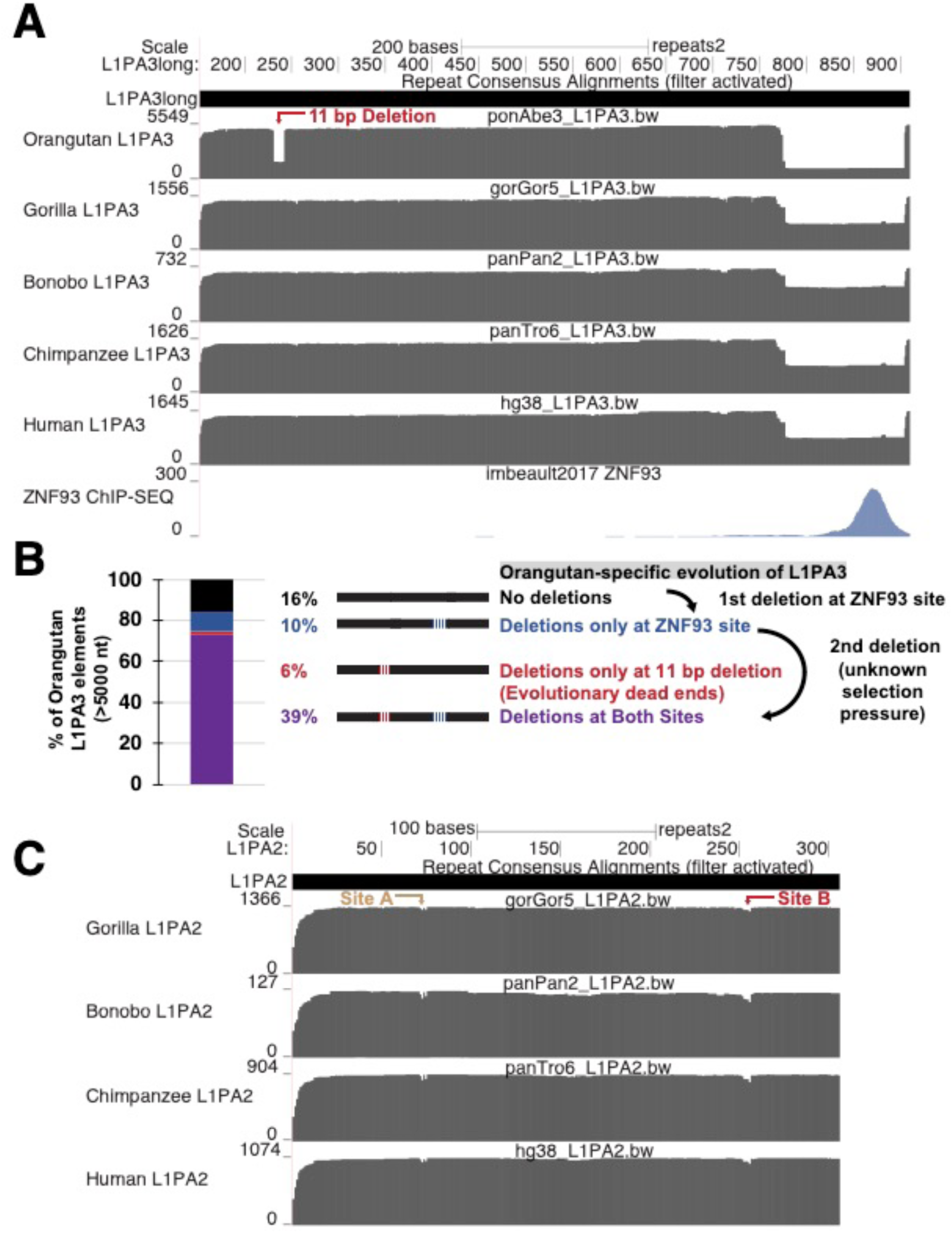
L1PA evolution in great apes. A) Coverage maps of L1PA3 demonstrate shared deletion of the ZNF93 binding site and an additional 11 bp deletion found only in orangutans. B) Analysis of the mutational pattern of orangutan elements suggests that the orangutan-specific mutation (red) occurred after ZNF93 evasion (blue) since this mutation is found almost exclusively in elements with the 129-bp deletion already. C) A) Coverage map of L1PA2 instances demonstrates no major changes across primates except for small deletions in an extreme 5’ region (Site A) and a region proximal to the orangutan deletion (Site B).

### No major deletions are visible in primate L1PA2 elements

Mapping of L1PA2 elements in gorilla, bonobo, chimp and human to the Repeat Browser reveals only minor changes between these relatively young elements. (Figure 6C) Although gorilla L1PA2 elements have greatly expanded compared to other primates, no significant gorilla-specific deletions are visible in our coverage plots; therefore the spread of gorilla elements may reflect the lack of a control factor that evolved in bonobo, chimpanzees and humans, or may reflect more subtle point mutations as we recently demonstrated for L1PA escape from ZNF649 control ^34^. Curiously, all four species show minor coverage drops in the area around nucleotide 250 (site B), a region that roughly corresponds to the deletion event observed in orangutan L1PA3 elements (Figure 6C). Although the deletion frequencies in primate L1PA2 are relatively low compared to the 11 bp L1PA3 orangutan deletion, this overall behavior is consistent with the model that this region is under adaptive selection - possibly to escape repression from a still unknown KZNF.

## DISCUSSION

The UCSC Repeat Browser provides an interactive and accessible environment to study repeat biology and side-steps the problem of mistakenly mapping reads to an incorrect locus by generating consensus representations of every repeat class, and focusing on how genome-wide datasets interact with repeat sequences independent of their genomic locus. Here we use this consensus-based approach to identify deletion events in repeats across species that suggest a model by which L1PA escape occurs differently across the phylogenetic tree of old world monkeys (Figure 7).

**FIGURE 7:**
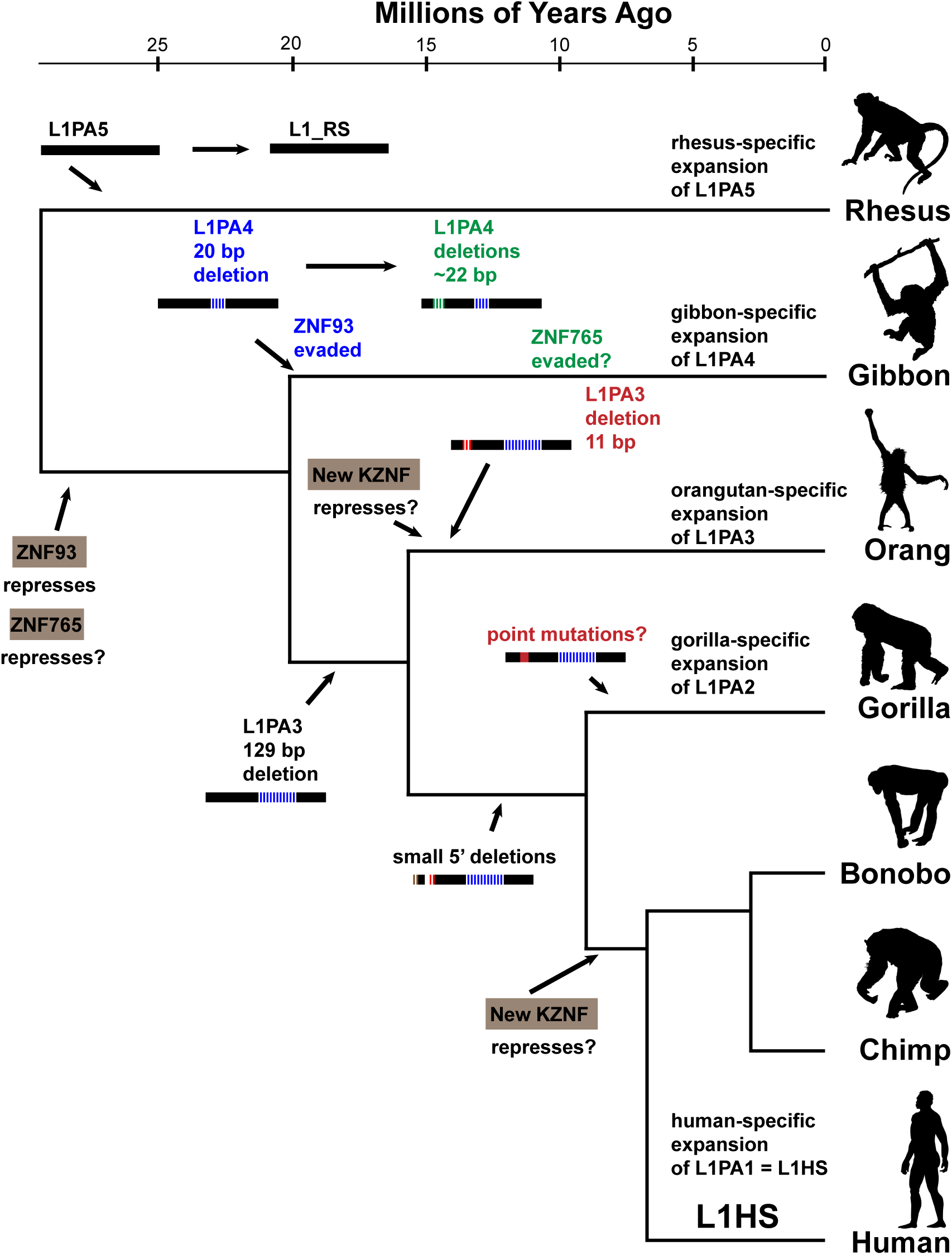
Model for L1PA evolution in different primate species. L1PA5 was active in the ancestor of human and rhesus, and expanded in a rhesus-specific fashion. ZNF93 evolved in the common ancestor of gibbons and humans (ape ancestor) to repress L1PA4 elements. In gibbons L1PA4 escaped with a small 20 bp deletion (blue); a second gibbon-specific deletion event (green) near the ZNF765 binding site led to gibbon-specific expansion of L1PA4. In great apes (human-orangutan ancestor) a 129 bp deletion (blue) in L1PA3 allowed ZNF93 evasion. In orang-utans (possibly in response to an orangutan specific KZNF) a new 11 bp deletion occurred and lead to orangutan-specific expansion of L1PA3. In gorillas, continued expansion of L1PA2 is not associated with deletions in the 5’UTR suggesting that this expansion is due either to lack of a chimp/bonobo/human repression factor or point mutations in gorilla L1PA2. A few gorilla, bonobo and human L1PA2 instances experience small deletions (brown and red); the red deletions are in a similar location to the orangutan L1PA3 deletion.

However, several caveats should be noted about Repeat Browser-based analyses. First, they rely entirely on RepeatMasker classifications (and in turn RepBase) and therefore depend on the quality of the annotations established in these collections. Second, the Repeat Browser uses its own consensus sequences to display genomic data, with these choices biased by length in order to ensure proper visualization, which can otherwise be problematic in regions where sequence is inserted. However, custom versions of the browser allow users to provide a custom consensus sequence. Indeed, we previously used this approach to create consensuses of L1PA3 subclasses (L1PA3long and L1PA3short (containing the ZNF93-related 129bp deletion)) when tracing an evolutionary arms race between ZNF93 and L1PA3 elements.^10^ Finally, the Repeat Browser and other consensus-based approaches risk diluting important, biologically relevant signal driven by a few instances of a repeat type that may affect the cell by virtue of their genomic location instead of their sequence. In these cases, the majority of instances in these families may generate no signal and produce an underwhelming “composite” Repeat Browser signal whereas an individual genomic locus may produce a strong, reproducible, and functionally relevant signal. Therefore, we recommend that Repeat Browser analysis be used in combination with existing genomic approaches for repeat analysis^9, 35–37^. Finally, the existence of the UCSC Repeat Browser as a complete “repeat genome collection” available for download should allow manipulation and utilization of repeat consensus sequences with a large set of existing, standard genomics tools, thereby enhancing the investigation of repeat sequence biology. We expect that the repeat community will make creative use of this tool beyond the workflows suggested here.

## CONCLUSIONS

The UCSC Repeat Browser provides a fully interactive environment, analogous to the UCSC Human Genome Browser, to study repeats. We show here that this environment provides an intuitive visualization tool for analysis and hypothesis-generation. For instance, here we use the Repeat Browser to demonstrate that sequence-specific deletions in repeats apparently driven by the activity of cellular repressors occurs independently in different species. The Repeat Browser is currently available at: http://bit.ly/repbrowser.

**Project name:** The UCSC Repeat Browser

**Project home page:** https://github.com/maximilianh/repeatBrowser

**Operating system(s):** Standard Web Browser

**Programming language:** Python, bash

**License:** Freely available for academic, nonprofit, and personal use.

**Any restrictions to use by non-academics:** Use of liftOver requires commercial license: http://genome.ucsc.edu/license

**Tutorial:** http://bit.ly/repbrowsertutorial

## FUNDING

This work was supported by EMBO ALTF 292-2011 and 4U41HG002371 to MH, F32GM125388 to JDF and 1R01HG010329 to SRS. DH is an investigator of the Howard Hughes Medical Institute.

## AUTHORS’ CONTRIBUTIONS

MH developed the concept for the Repeat Browser with input from all other authors. JDF developed the Repeat Browser tutorial and materials for general release. JDF and AZ analyzed KZNF and repeat data. MH, JDF, SRS, WJK and DH conceived of the idea and contributed to the Repeat Browser’s design. JDF, SRS and MH wrote the manuscript.

## ACKNOWLEDGEMENTS

We thank A. Smit, R. Hubley, and A. Ewing for helpful discussions about repeat consensus choice and annotations. We thank J. Armstrong, F. Jacobs and D. Greenberg and all members of the Haussler lab for helpful comments and discussion.

## Notes

#### Summary of Updates

This version updates the genome assemblies analyzed, clarifies analyses and provides an updated version of the Repeat Browser that is both hg19 and hg38 compatible.

https://genome.ucsc.edu/s/jdf2001/courtyard_browser

## REFERENCES

1. Slotkin, R. K. The case for not masking away repetitive DNA. Mob. DNA 9, 15 (2018).

2. Chuong, E. B., Elde, N. C. & Feschotte, C. Regulatory activities of transposable elements: from conflicts to benefits. Nat. Rev. Genet. (2016). doi:10.1038/nrg.2016.139

3. Pastuzyn, E. D. et al. The Neuronal Gene Arc Encodes a Repurposed Retrotransposon Gag Protein that Mediates Intercellular RNA Transfer. Cell 173, 275 (2018).

4. Ding, Y., Berrocal, A., Morita, T., Longden, K. D. & Stern, D. L. Natural courtship song variation caused by an intronic retroelement in an ion channel gene. Nature 536, 329–332 (2016).

5. Tam, O. H. et al. Postmortem Cortex Samples Identify Distinct Molecular Subtypes of ALS: Retrotransposon Activation, Oxidative Stress, and Activated Glia. Cell Rep. 29, 1164–1177.e5 (2019).

6. Bao, W., Kojima, K. K. & Kohany, O. Repbase Update, a database of repetitive elements in eukaryotic genomes. Mob. DNA 6, 11 (2015).

7. Smit, A., Hubley, R. & Green, P. RepeatMasker Open-4.0. http://www.repeatmasker.org

8. Jurka, J., Walichiewicz, J. & Milosavljevic, A. Prototypic sequences for human repetitive DNA. J. Mol. Evol. 35, 286–291 (1992).

9. Goerner-Potvin, P. & Bourque, G. Computational tools to unmask transposable elements. Nat. Rev. Genet. 1 (2018). doi:10.1038/s41576-018-0050-x

10. Jacobs, F. M. J. et al. An evolutionary arms race between KRAB zinc-finger genes ZNF91/93 and SVA/L1 retrotransposons. Nature 516, 242–5 (2014).

11. Sun, X. et al. Transcription factor profiling reveals molecular choreography and key regulators of human retrotransposon expression. Proc. Natl. Acad. Sci. U. S. A. 115, E5526–E5535 (2018).

12. Edgar, R. C. MUSCLE: multiple sequence alignment with high accuracy and high throughput. Nucleic Acids Res. 32, 1792–7 (2004).

13. Kent, W. J. et al. The Human Genome Browser at UCSC. Genome Res. 12, 996–1006 (2002).

14. Haeussler, M. et al. The UCSC Genome Browser database: 2019 update. Nucleic Acids Res. 47, D853–D858 (2019).

15. Kent, W. J. BLAT---The BLAST-Like Alignment Tool. Genome Res. 12, 656–664 (2002).

16. Benson, G. Tandem repeats finder: a program to analyze DNA sequences. Nucleic Acids Res. 27, 573–80 (1999).

17. Rice, P., Longden, I. & Bleasby, A. EMBOSS: the European Molecular Biology Open Software Suite. Trends Genet. 16, 276–7 (2000).

18. Kohany, O., Gentles, A. J., Hankus, L. & Jurka, J. Annotation, submission and screening of repetitive elements in Repbase: RepbaseSubmitter and Censor. BMC Bioinformatics 7, 474 (2006).

19. Altschul, S. F., Gish, W., Miller, W., Myers, E. W. & Lipman, D. J. Basic local alignment search tool. J. Mol. Biol. 215, 403–410 (1990).

20. Harrow, J. et al. GENCODE: the reference human genome annotation for The ENCODE Project. Genome Res. 22, 1760–74 (2012).

21. Hsu, F. et al. The UCSC Known Genes. Bioinformatics 22, 1036–1046 (2006).

22. ENCODE Project Consortium. An integrated encyclopedia of DNA elements in the human genome. Nature 489, 57–74 (2012).

23. Imbeault, M., Helleboid, P.-Y. & Trono, D. KRAB zinc-finger proteins contribute to the evolution of gene regulatory networks. Nature 543, 550–554 (2017).

24. Najafabadi, H. S. et al. C2H2 zinc finger proteins greatly expand the human regulatory lexicon. Nat. Biotechnol. 33, 555–62 (2015).

25. Thomas, J. H. & Schneider, S. Coevolution of retroelements and tandem zinc finger genes. Genome Res. 21, 1800–12 (2011).

26. Leinonen, R., Sugawara, H., Shumway, M. & International Nucleotide Sequence Database Collaboration. The sequence read archive. Nucleic Acids Res. 39, D19–21 (2011).

27. Langmead, B. & Salzberg, S. L. Fast gapped-read alignment with Bowtie 2. Nat. Methods 9, 357–359 (2012).

28. Zhang, Y. et al. Model-based analysis of ChIP-Seq (MACS). Genome Biol. 9, R137 (2008).

29. Khan, H., Smit, A. & Boissinot, S. Molecular evolution and tempo of amplification of human LINE-1 retrotransposons since the origin of primates. Genome Res. 16, 78–87 (2006).

30. Konkel, M. K., Walker, J. A. & Batzer, M. A. LINEs and SINEs of primate evolution. Evol. Anthropol. 19, 236–249 (2010).

31. Han, K. et al. Mobile DNA in Old World monkeys: A glimpse through the rhesus macaque genome. Science (80-.). 316, 238–240 (2007).

32. Schmitges, F. W. et al. Multiparameter functional diversity of human C2H2 zinc finger proteins. Genome Res. 26, 1742–1752 (2016).

33. Larson, P. A. et al. Spliced integrated retrotransposed element (SpIRE) formation in the human genome. PLoS Biol. 16, (2018).

34. Fernandes, J. D. et al. KRAB Zinc Finger Proteins coordinate across evolutionary time scales to battle retroelements. bioRxiv 429563 (2018). doi:10.1101/429563

35. Jeong, H. H., Yalamanchili, H. K., Guo, C., Shulman, J. M. & Liu, Z. An ultra-fast and scalable quantification pipeline for transposable elements from next generation sequencing data. in Pacific Symposium on Biocomputing 0, 168–179 (World Scientific Publishing Co. Pte Ltd, 2018).

36. Jin, Y., Tam, O. H., Paniagua, E. & Hammell, M. TEtranscripts: A package for including transposable elements in differential expression analysis of RNA-seq datasets. Bioinformatics 31, 3593–3599 (2015).

37. Kong, Y. et al. Transposable element expression in tumors is associated with immune infiltration and increased antigenicity. Nat. Commun. 10, 5228 (2019).

